# A Generalizable Tension Sensor Platform for Mechanotherapeutic Discovery

**DOI:** 10.1101/2025.05.15.653325

**Authors:** Matthew R. Pawlak, Andrew M. Baldys, Matthew C. McMahon, Frank M. Cichocki, Wendy R. Gordon

## Abstract

Mechanical forces are critical regulators of cellular function, and their modulation represents a promising therapeutic strategy across a range of diseases, including cancer and fibrosis. DNA-based molecular tension sensors (MTSs) have emerged as powerful tools for detecting receptor-specific cellular forces but remain limited by susceptibility to nuclease degradation and constrained ligand compatibility. Here, we outline these barriers to broader adoption and demonstrate how integrating established stabilization strategies effectively mitigates nuclease sensitivity. In addition, we introduce an engineered protein that covalently couples ligands to DNA in a modular, receptor-agnostic manner. Together, these innovations enable robust, nearly universal deployment of DNA-based MTSs across diverse experimental contexts and target proteins. We apply this enhanced platform to profile the mechanobiological effects of force-modulating drugs in both immortalized and primary cell lines. Unlike indirect or context-limited methods, this approach delivers direct, quantitative readouts of drug-induced changes in mechanical force transmission, offering a scalable path toward personalized mechanotherapeutic screening.

## Introduction

Mechanical forces are fundamental regulators of cellular function, influencing processes such as motility^1^, adhesion ^2^, differentiation^3^, and chromatin accessibility ^4^. These forces are transmitted through the cytoskeleton and surface receptors to the surrounding extracellular matrix (ECM), and/or neighboring cells. Dysregulated mechanobiology contributes to numerous diseases, including cancer ^5^, fibrosis ^6^, and immune disorders ^7^. Consequently, the ability to quantify and manipulate cellular mechanics presents both a diagnostic opportunity and a therapeutic avenue.

Mechanotherapeutics, which aim to modulate cellular mechanics, represent a promising but underdeveloped class of therapies^8^. Integrins, a diverse family of adhesion receptors, are particularly attractive targets due to their role in force transmission and mechanosignaling^9^. Integrin-targeting drugs have shown clinical success in conditions such as multiple sclerosis and thrombosis ^10^, yet their widespread therapeutic adoption has been hindered by an incomplete understanding of integrin activation mechanisms ^11,12^ and limited functional screening methods. Current adhesion assays lack precision and fail to capture the full spectrum of integrin activity^13^, while luminescence-based approaches require engineered integrins ^14^, restricting their applicability to primary cells and personalized medicine.

Molecular tension sensors (MTSs) offer a powerful alternative for assessing force generation by cells because they probe specific cell-to-environment interactions, typically via fluorescence-based readouts^15^. A widely used subclass of MTSs, tension gauge tethers (TGTs), consist of DNA duplexes which rupture upon transmission of a tunable critical force (Fig. S1), enabling quantitative measurement of receptor-mediated forces^16,17^. Our recent advancement, Rupture and Deliver TGTs (RAD-TGTs), enhances this approach by tagging force-exerting cells with fluorescent payloads in proportion to their cellular forces, enabling high-throughput analysis through flow cytometry ^18^. Using this strategy, we demonstrated the ability to rapidly measure mechanical profiles of thousands of cells, providing a relative quantification of mechanical phenotype, or mechanotype. In the same work we also introduced an alternative strategy to append ligands to TGTs using HUH endonucleases (HUH), a family of small nucleases which bind to and form a covalent adduct with DNA in a site-specific manner with high efficacy and no further purification ^19,20^. HUHs were able to retain function when fused to ligands including the pan-RGD binding integrin binder echistatin^21^, demonstrating a facile method to append ligands to TGTs.

Despite their potential, MTSs, including TGTs, remain underutilized beyond the labs that develop them. Expanding their adoption requires addressing key limitations, including the susceptibility of DNA-based MTSs to nucleases^22^ and the limited availability of easily conjugated ligands beyond integrins ^23^. While integrins serve as an excellent model system due to their direct force-dependent activation, broadening the ligand repertoire is critical to making NATS (Nucleic Acid Tension Sensors) applicable to a wider range of mechanotherapeutic targets. The introduction of nuclease-resistant alternative nucleic acids, such as peptide nucleic acid (PNA):DNA hybrids ^24^, provides a potential solution, though clear guidelines for their use are lacking.

Outside of RAD-TGTs, a growing assortment of high-throughput NATS have emerged, each with their own benefit. In particular, the Salaita group has developed high-throughput assessments of NATS utilizing flow cytometry (µTS^25^ and TaCT^26^) and plate reader-based assays (MCATS^27^). These assays provide the first scalable methods to assess cellular mechanics at a single-molecule level. Such methods stand to expand not only the understanding of mechanobiology when paired with state-of-the-art -omics analysis, but the development of mechanotherapeutics. High-throughput NATS possess the unique ability to assess the direct effect of mechanotherapeutics on their intended targets. For example, integrin activity is directly related to the mechanical force transmitted; high-throughput NATS can be used to assess how different putative modulators alter activity at varying concentrations, yielding what may be considered a “mechano-kinetic” analysis. Furthermore, NATS do not require engineered cell lines and may be applicable to a wide variety of cell types. The barriers laid out above limit the current utility of NATS, and strategies to overcome said challenges must be integrated into experimental practices for NATS to achieve their potential. In this work, we address these barriers by establishing a universal NATS platform compatible with diverse cell lines and therapeutic targets. We demonstrate strategies to mitigate nuclease degradation and introduce a novel method for covalently linking IgG-containing proteins to DNA structures using engineered Protein G^28^ (PG)-HUH fusion. These advancements are not limited to integrins but provide a broadly applicable strategy for studying mechanotransduction across multiple receptor types. By integrating these developments into RAD-TGTs, we provide a robust screening platform for mechanotherapeutics which can be used to profile how these treatments alter a cell’s mechanical phenotype (mechanotype). We demonstrate this function by analyzing the effect of integrin modulatory agents on both immortalized cell lines and primary immune cells. This work lays the foundation for expanding MTS applications beyond specialized mechanobiology labs, making them accessible for broader use in drug development and mechanomedicine.

## Results and Discussion

### Nuclease Activity in DNA-Based Nucleic Acid Tension Sensors

A critical limitation of DNA-based molecular tension sensors (MTSs) is their susceptibility to degradation by nucleases, which can originate from serum components in culture media ^29^ or from cellular sources such as secreted ^30^ or membrane-bound nucleases (e.g., DNase X, which is highly expressed in many cancers ^31^). Nuclease activity introduces variability into DNA-based MTS assays by degrading probes unpredictably, leading to inconsistent ligand presentation and volatile readouts. Despite its widespread impact, nuclease activity remains underreported in MTS studies, and strategies to mitigate this challenge are still emerging. The Wang group has been at the forefront of these efforts, introducing both a surface nuclease sensor (SNS) ^32^ to visualize nuclease activity (Fig. S2) and DNA:PNA hybrids as a means to enhance nuclease resistance while maintaining tunable rupture forces^24^.

To systematically characterize how nuclease activity manifests under our typical experimental settings, we first employed SNS probes to visualize nuclease activity across different conditions. First, we assessed the contribution of exogenous nucleases by plating U-251 cells (low nuclease activity ^18^) on SNS-coated surfaces in serum-free and full-serum media. Under serum-free conditions, SNS fluorescence was negligible, whereas in serum-containing media, a robust SNS signal was observed throughout the imaging field (Figure **1A,B**). This suggests that exogenous nucleases degrade DNA probes indiscriminately across the experimental surface, potentially interfering with MTS-based readouts.

**Figure 1.**
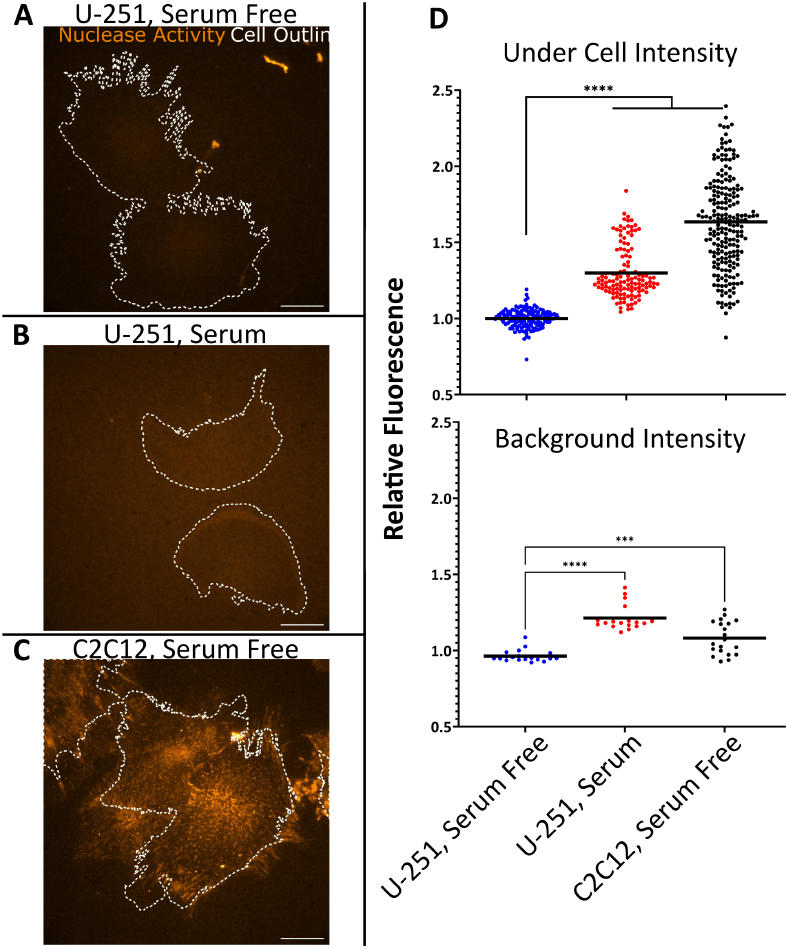
Nuclease Activity is Persistent Throughout Experiments. A–C) Representative images of nuclease activity across different conditions. Orange intensity represents SNS-detected nuclease activity; white dotted lines show cell outlines. Scale bar = 20 µm. A) U-251 cells in serum-free media. B) U-251 cells in serum-containing media. C) C2C12 cells in serum-free media. D) Quantification of activity: Median fluorescence intensity under cells (top) vs background (bottom), normalized to U-251 serum-free under-cell intensity. Data are means of replicate-normalized values analyzed via ordinary one-way ANOVA. **** *P <* 0.0001, *** *P* = 0.0001.

To evaluate the impact of endogenous nucleases, we plated C2C12 cells (high extracellular nuclease activity ^33^) on SNS surfaces in serum-free media. These cells exhibited strong, localized SNS signals directly beneath their adhesion footprints, consistent with prior reports of DNase X activity (Figure **1C**) ^31,32^. Quantification of SNS intensity under adhered cells versus background regions confirmed significant differences in nuclease activity between conditions (Figure **1D**). These findings highlight the challenges posed by both exogenous and endogenous nucleases in DNA-based MTS applications. Furthermore, alternative nucleic acids are more costly than traditional DNA oligonucleotides, thus all experimental components must be assessed for nuclease activity before backbone composition is decided.

### DNA:PNA Hybrids Enhance Fidelity of High-Throughput Mechanotyping

Given the limitations imposed by nuclease activity on DNA-based RAD-TGTs (**Figure 2A-B**), we investigated whether DNA:PNA hybrid duplexes could provide a nuclease-resistant alternative suitable for high-throughput mechanotyping. To evaluate this, we performed RAD-TGT assays using DNA or DNA:PNA duplexes on low (U-251) and high (C2C12) nuclease activity cells, with and without pharmacological perturbation using the ROCK inhibitor Y-27632. For this experiment, cells were plated onto surfaces exclusively containing the high rupture force shearing conformation TGTs, as prior reports demonstrate Y-27632’s effects on TGT rupture were most robust in the shearing conformation ^34^. To serve as ligand, TGTs were reacted with HUH alone to serve as a negative control or with HUH-echistatin to specifically target the forces transmitted through RGD-binding integrins.

**Figure 2.**
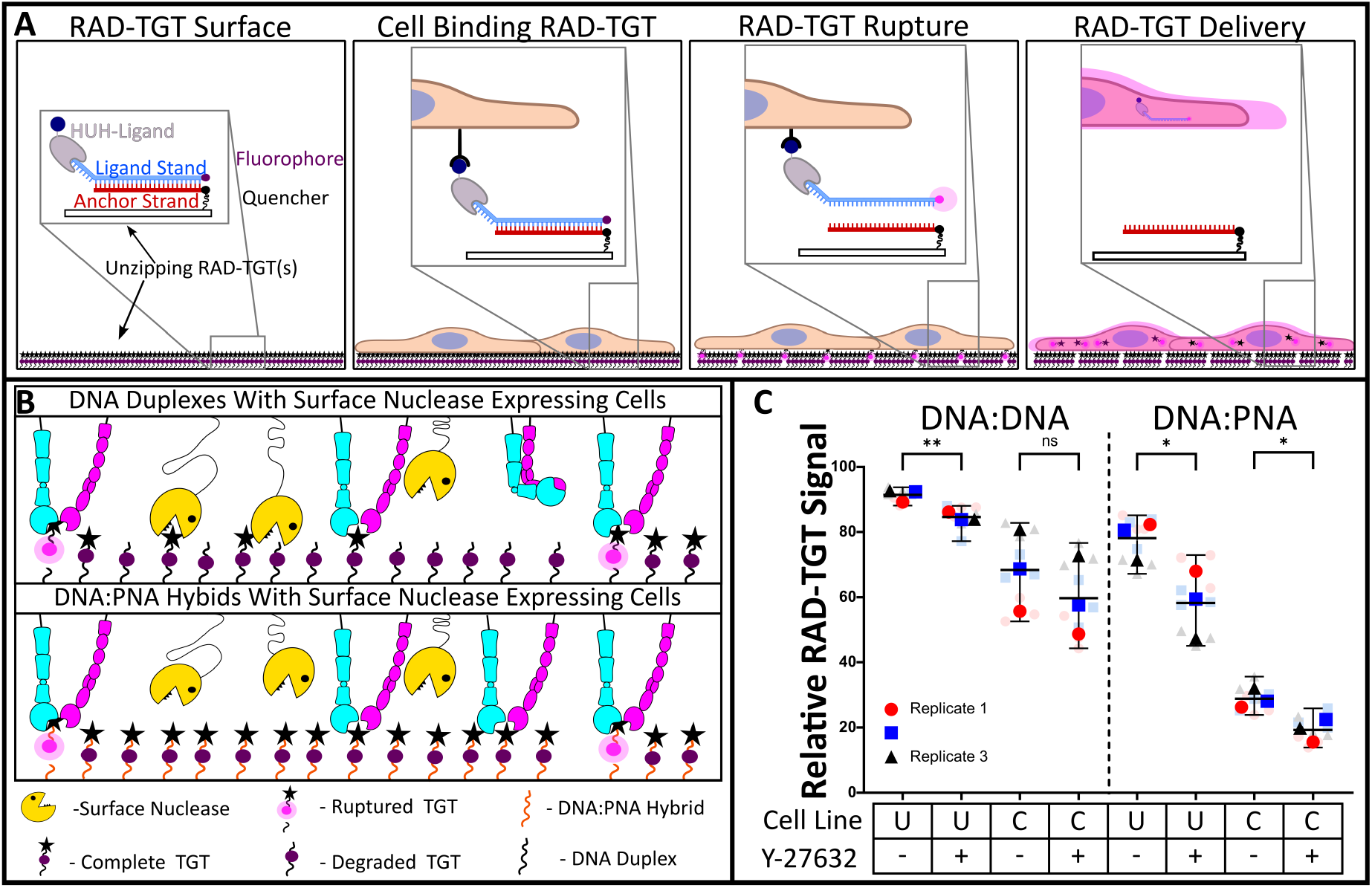
Overview of RAD-TGT Mechanism and Nuclease Effects. (A) Cartoon schematic of RAD-TGT function. Surfaces coated with TGTs are incubated with cells, which bind to ligands presented by TGTs. If sufficient force is transmitted through the TGT, the duplex ruptures, separating the fluorophore–quencher pair. The fluorophore-containing strand is delivered to the cell, increasing cellular fluorescence. (B) Cartoon illustrating the effect of surface nucleases on NATS composed of DNA or DNA:PNA hybrids. DNA NATS (top) are susceptible to degradation by surface nucleases, leading to a local decrease in ligand availability (cyan and pink dimers), which introduces artifacts. DNA:PNA hybrid NATS resist degradation, improving data fidelity. (C) RAD-TGT results in low-nuclease (U-251) and high-nuclease (C2C12) cell lines, with or without drug treatment, on either DNA:DNA duplexes (left) or DNA:PNA hybrids (right). RAD-TGT signal represents the percentage of the population exceeding the 99th percentile of the respective no-ligand control. Each replicate is represented by a distinct color and shape; solid colors indicate the average of two technical replicates, while opaque symbols represent biological replicates. Cell lines are denoted as U (U-251) or C (C2C12); Y-27632-treated samples are marked with +, vehicle controls with –. Means of technical replicates were compared using unpaired *t*-tests (*N* = 3): ns = not significant, \**P <* 0.05, \*\**P <* 0.01. Error bars represent standard deviation.

We hypothesized that DNA-based RAD-TGTs would reliably detect force reductions following drug treatment in low-nuclease activity cells but would fail to do so in high-nuclease activity cells due to probe degradation. In contrast, we expected that DNA:PNA hybrids would retain sensitivity to force changes regardless of nuclease activity. As anticipated, U-251 cells exhibited significant reductions in RAD-TGT signal following Y-27632 treatment on both DNA and DNA:PNA duplexes. Notably, the absolute RAD-TGT signal was lower for DNA:PNA hybrids due to their greater mechanical stability; however, when assessing population shifts relative to respective negative controls, both probe types yielded statistically significant fold reductions in signal (**Figure 2C**). Additionally, assessing the relative fold change of median fluorescent intensity of the population supported these findings (**Figure S4**). In contrast, C2C12 cells showed no significant change in RAD-TGT signal following drug treatment when analyzed using DNA-based probes, confirming that nuclease activity masked force-dependent changes. Strikingly, when analyzed on DNA:PNA hybrid probes, C2C12 cells exhibited a significant reduction in RAD-TGT signal following Y-27632 treatment, similar to U-251 cells (**Figure 2D**).

These results demonstrate that nucleases can substantially limit the utility of DNA-based MTSs in high-nuclease environments, but these limitations can be effectively overcome with nuclease-resistant oligonucleotides. The successful application of DNA:PNA hybrids in RAD-TGT assays supports their broader adoption in mechanotyping studies, particularly for high-throughput screening applications in cell lines that were previously incompatible with DNA-based probes.

### Covalent Protein-G Expands Ligand Repertoire

The second major limitation of NATS we wanted to address is the restricted repertoire of available ligands. Most studies utilizing NATS have used the ligand cyclic RGD owing to its ability to bind to one-third of all integrin heterodimers ^21^ and its robust thermostability, allowing it to function after high-temperature DNA annealing ^35^. Alternative peptide sequences such as LDVP can allow targeting of α4β1 integrins^36^, yet this is just a fraction of all bona fide and putative adhesion and force-transmitting receptors. Furthermore, most strategies to append the ligand to DNA-based sensors require chemical synthesis and purification, which once again proves to be an impediment to adaptation.

In 2016, Wang et al. proposed the use of Protein G (PG), a small Fc-binding protein, to functionalize TGTs with Fc-tagged ligands or antibodies^23^. This strategy provides a “plug-and-play” approach that simplifies and expands ligand attachment, but concerns remain regarding the mechanical stability of the PG-Fc interaction. Previously, we utilized HUH-ligand fusions to append ligands of interest to NATS and initially saw this as an accessible method to expand the ligand repertoire. Although successful, this method required expression and purification of each potential ligand of interest. We sought to streamline this method by fusing PG with a HUH. In theory, this resulting protein would be able to covalently bind the TGT while also binding an Fc-containing ligand, yielding a universal ligand attachment strategy that circumvents the need for chemical synthesis.

We first fused the first IgG-binding domain of PG to the N-terminus of a HUH. This construct retained the ability to covalently bind DNA (**Figure S6**) and was used to attach an anti-β1 integrin antibody (K20) to both unzipping and shearing TGTs. When tested with U-251 cells, this system supported some, albeit minimal, cell adhesion, but cell density remained low and individual cells struggled to spread (**Figure 3B**). To determine whether this behavior resulted from using an antibody rather than a physiological ligand—or from the nature of the PG attachment—we expressed a direct HUH-K20 fusion protein (see preprint) and repeated the adhesion assay. Surprisingly, cells adhered and spread robustly, suggesting that the PG-Fc interaction was responsible for the weaker adhesion (**Figure 3A**).

**Figure 3.**
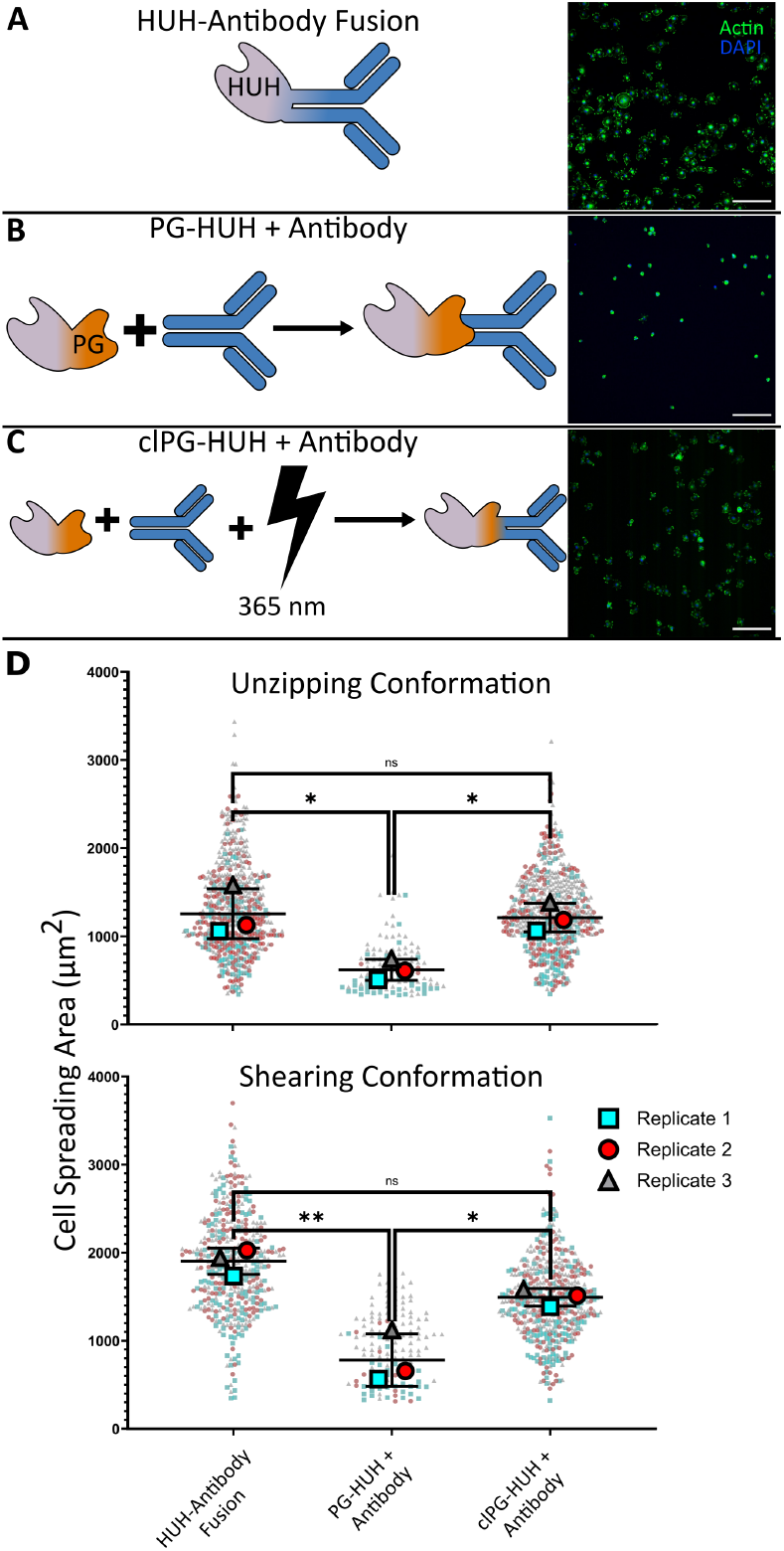
Figure 3: Covalent Protein-G Expands Lig- and Repertoire. (A–C) Cartoon schematics and representative images showing three methods for appending antibodies to TGTs using HUH fusions. (A) Direct fusion of the HUH enzyme to the antibody of interest. (B) HUH-Protein G (PG) fusion, which binds non-covalently to Fc-containing proteins. (C) clPG-HUH fusion, which covalently binds antibodies following incubation and exposure to 365 nm light. (D) Quantification of cell spreading on TGTs presenting antibodies via each method: direct fusion, PG-HUH, and clPG-HUH (left to right). Top: unzipping TGTs; bottom: shearing TGTs. Data are shown as a superplot of three biological replicates. Large opaque points represent replicate means; small transparent points are individual cells (only singular, non-touching cells included; minimum *n* = 20 per replicate). Replicate means were compared using ordinary one-way ANOVA (*N* = 3): \*\**P <* 0.01, \**P <* 0.05, ns = not significant Scale Bar = 250 µm.

We hypothesized that, although PG is mechanically stronger than the DNA duplex, its non-covalent Fc interaction ruptures before DNA rupture due to transient binding or applied forces ^37,38^. This premature ligand detachment would reduce effective ligand density, thereby impairing cell adhesion and spreading. To address this, we developed a modified PG-HUH construct incorporating benzoyl-phenylalanine, an unnatural photocrosslinking amino acid, at the IgG-binding interface. This modification enables covalent crosslinking of PG to Fc-tagged ligands upon 365 nm light activation and has previously been used to covalently link proteins of interest to antibodies^28,39^. We aptly named the resulting construct crosslinking PG-HUH (clPG-HUH). To confirm crosslinking, clPG-HUH was incubated with K20 and exposed to blue light. This sample was then analyzed via SDS-PAGE, and the resulting gels showed an apparent gain of mass, confirming successful crosslinking (**Figure S7**). In a comparable manner, we confirmed that HUH retained DNA binding capability, yet longer exposure led to decreased HUH efficacy. Comparing HUH efficacy to crosslinking efficacy, we determined a 20-minute light exposure yielded the most crosslinking while retaining HUH function (**Figures S8–S9**).

When applied in the K20 adhesion assays, this clPG-HUH restored cell adhesion and spreading to levels comparable to direct ligand fusion (**Figure 3C**). Quantification of the spread area of single cells confirmed visual assessment of the different ligand attachment methods (**Figure 3D**). For cells plated on unzipping TGTs, both the HUH-K20 and clPG-HUH had an average spreading area of ∼1200 µm^2^, yet the non-crosslinking PG-HUH had a statistically significant decrease to ∼600 µm^2^. For shearing TGTs, the statistical insignificance between spreading area of the fusion and clPG-HUH was maintained, as well as the statistically significant decrease between these and the non-crosslinking PG-HUH. Average spreading areas of the HUH-K20 fusion, clPG-HUH, and non-crosslinking PG-HUH were ∼1900 µm^2^, ∼1500 µm^2^, and ∼800 µm^2^, respectively. Although not significant, the decrease between the HUH fusion and clPG-HUH was still apparent, which we attributed to either incomplete crosslinking or slightly decreased ligand abundance caused by decreased HUH function following blue light exposure. These results highlight that while PG-based TGTs provide a versatile ligand selection platform, fidelity of the non-covalent ligand attachment is a critical factor that may otherwise confound data interpretation. By integrating covalent crosslinking, this system allows any Fc-containing ligand to be used with DNA MTSs while maintaining high mechanical stability and reproducibility.

### Assessing the Efficacy of Mechanotherapeutics

Changes in cellular mechanotype often accompany disease progression, prompting interest in pharmacological strategies to restore healthy mechanics and function. However, few mechanobiology assays exist that are compatible with high-throughput drug screening. With the improvements described above, NATS—particularly RAD-TGTs—are well-suited for this role. They are broadly applicable across cell types, media conditions, ligand choices, and nuclease environments. Importantly, the RAD-TGT assay is accessible to molecular biology labs, as all components are either commercially available or easily recombinantly expressed. We hypothesized that RAD-TGTs could serve as a universal platform to assess how drugs alter cell adhesion dynamics (specifically receptor abundance and mechanical engagement), providing a direct readout of mechanotherapeutic efficacy (**Figure 4**).

**Figure 4.**
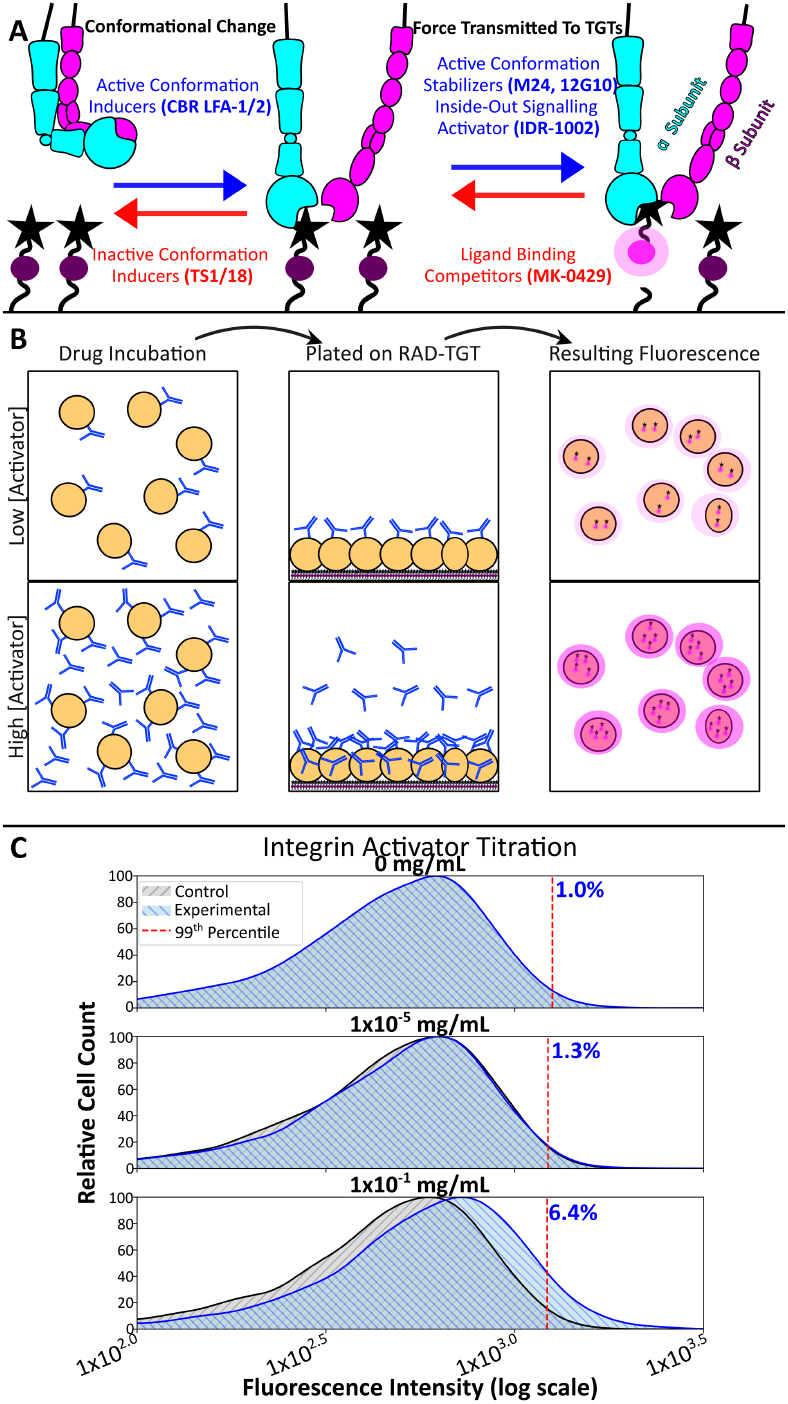
Principle of High-Throughput NATS Assessing Mechanotherapeutics. A) How integrin targeting therapeutics alter mechanical force transmission. Integrins (cyan and pink dimers) are in a resting state and fail to engage RAD-TGTs(star and lavender circle), conformation inducing therapeutics (activators in blue, inhibitors in red) can lead to conformational change to the active conformation or return to the inactive state. Once engaged with the RAD-TGT, conformational stabilizers or inside out activators permit the exchange of mechanical force resulting in rupture and gain of fluorescence. This process can be inhibited via ligand binding competitors. B) Theoretical principle behind assessing mechanokinetics via RAD-TGTs. Cells (yellow circles) are treated with varying concentrations of activating therapeutic (blue antibody). Following incubation cells are plated on RAD-TGT coated surfaces. Engagement with and subsequent rupture of RAD-TGTs result in proportional increases of fluorescence. C) Representative data of a titration of activator, as concentration increases the population experiences an increase in fluorescence relative to respective vehicle control.

To evaluate this, we analyzed how LFA-1 (a primary integrin in immune cells) modulating drugs alter the mechanotype of primary human natural killer (NK) cells. This system was selected because the role of mechanical forces in immune cell function (a field aptly named immunomechanics) has rapidly risen to prominence in the past 5 years and with that rise so has the interest in pharmacological modulation of immunomechanics ^40–42^.Assessing donor-collected cells directly also provides the foundation for personalized mechanomedicine. One can envision how directly assessing a patient’s immune cells’ response to mechanotherapeutics can provide invaluable insights towards crafting bespoke therapeutic strategies.

To assess the validity of this strategy, the incorporation of the advancements in this work is necessary. NK cells require serum-containing media, and thus nuclease-resistant DNA:PNA hybrids are used.

Furthermore, our desired therapeutic target (LFA-1) does not bind RGD-presenting ligands but rather binds ICAM-1. Fortunately, ICAM-1 Fc fusion proteins are commercially available and can be appended to our DNA:PNA tethers via clPG-HUH. We sought to characterize separate classes of LFA-1 targeting drugs, in particular inhibitors, activators, and reporters. To achieve this, we settled on the following antibodies: TS1/18 (an inhibitor which prevents LFA-1 from binding ligands) ^43^, CBR LFA-1/2 (an activator) ^44^, and M24 (activation reporter) ^45^ (**Fig 4A**).

To quantify any changes in LFA-1 force signature, the 99th percentile of the RAD-TGT signal was identified for each respective isotype control, and the percent of each corresponding experimental population was calculated. This analysis generated dose-dependent responses by every therapeutic similar to effective concentration (EC_50_) or inhibitory concentration (IC_50_) curves dependent on the modulatory function of the treatment. The inhibitor, TS1/18, produced a robust decrease of signal as concentration increased, confirming both that TS1/18 is indeed an inhibitor of LFA-1 mediated forces and that this assay could be used to assess therapeutic-induced changes of mechanotype. We fit an IC_50_ curve to this response to quantify the mechano-IC_50_ and found a value of 0.5 ± 0.2 ng/ml (3 nM), indicating a robust mechanoresponse to this treatment. The known activator, CBR LFA-1/2, and the activation reporter, M24, generated similar curves, which were the inverse of TS1/18, indicating both behaved as mechano-activators (**Fig 5A-B**). Both mechano-activators demonstrated unique behavior; as treatment concentration increased, so did RAD-TGT signal until a relative peak was achieved and RAD-TGT signal began to decrease (referred going forward as a mechano-maxima). CBR LFA-1/2 achieved both a lower mechano-EC_50_ and lower mechano-maxima than M24, with EC_50_ values of 0.1 ± 0.1 ng/ml (0.6 nM) and 1 ± 0.4 (6 nM) and mechano-maxima at 1 ng/ml and 10 ng/ml for CBR LFA-1/2 and M24, respectively (**Fig 5B**).

**Figure 5.**
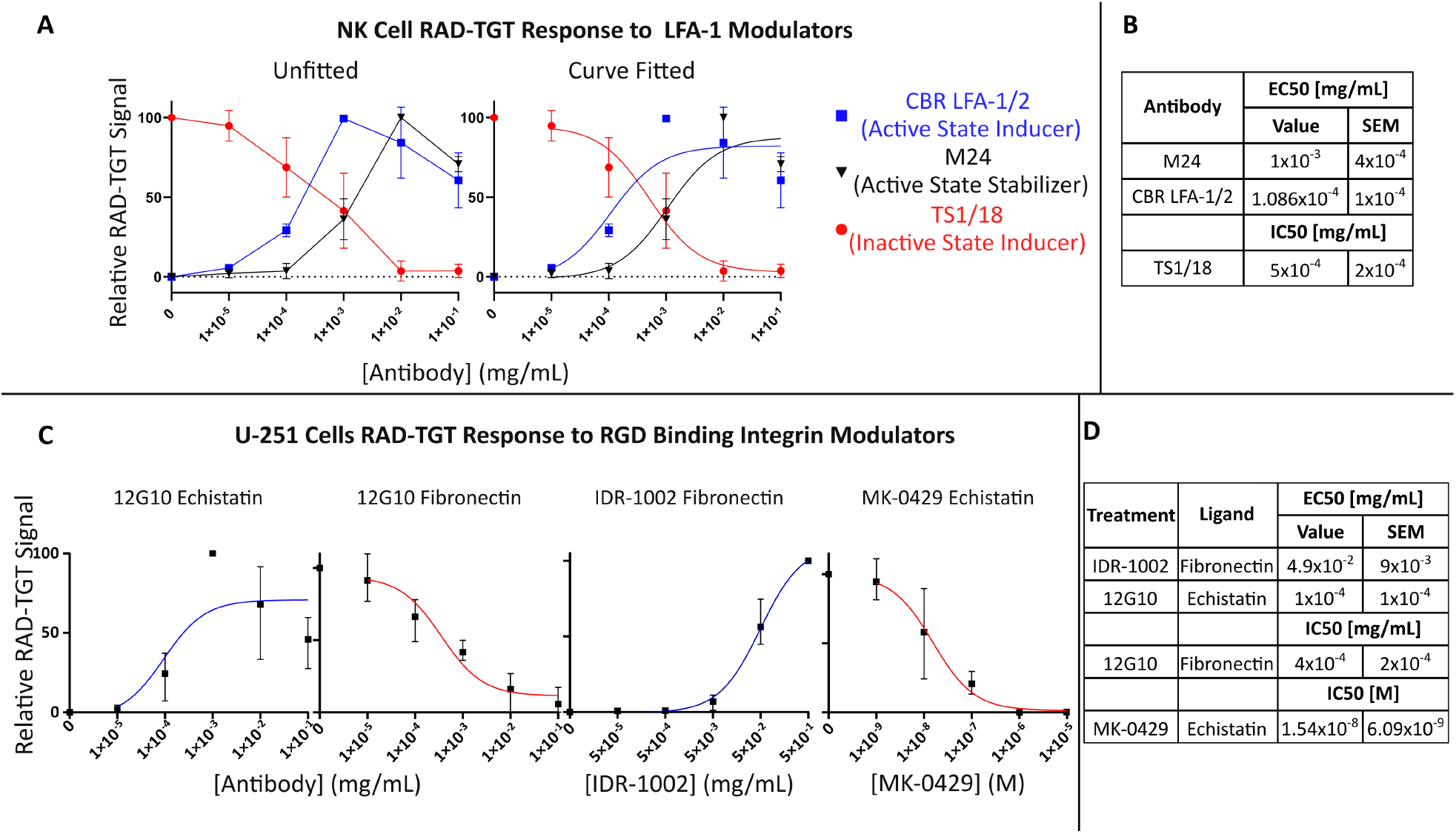
Mechanokinetic Assessment of Integrin Modulators. (A–B) Quantification of the effects of LFA-1–modulating antibodies on NK cells using unzipping RAD-TGTs presenting ICAM-1. (A) RAD-TGT signal generated across a titration series of CBR LFA-1/2 (blue squares), M24 (black triangles), and TS1/18 (red circles). Data are presented as the mean percentage of the cell population exceeding the 99th percentile of the corresponding vehicle control, normalized from 0 to 100 using the lowest and highest concentrations of treatment. Each point represents the mean of three biological replicates, each composed of technical duplicates containing *>*3,000 cells. The left panel shows data points connected to adjacent titration steps; the right panel displays fitted dose-response curves using either an agonist or inhibitor three-parameter model. Error bars represent standard deviation. (B) Summary of EC_50_ or IC_50_ values derived from the fits in (A), including the mean and standard error. (C–D) Mechanokinetic assessment of RGD-binding integrin-targeting drugs in U-251 cells using unzipping RAD-TGTs. (C) Dose-response curves for U-251 cells treated with the indicated drug, with ligand identity listed in each title. Data are presented as in (A), representing three independent experiments with ¿1,000 cells each. (D) Summary of EC_50_ or IC_50_ values for each condition in (C), including the mean and standard error.

These results offer insight into both the nature of the therapeutics and the assay itself. Both CBR LFA-1/2 and TS1/18 were able to recapitulate their reported function as activator or inhibitor, highlighting their impressive function with low to sub-nanomolar mechano-EC_50_/IC_50_ values, respectively. When directly comparing the bona-fide (CBR LFA-1/2) and putative (M24) activators, CBR LFA-1/2 demonstrated greater activating abilities compared to M24, consistently demonstrating a mechano-EC_50_ an order of magnitude lower than M24. Despite this lesser capability, M24 still had notable capability to function as an activator, demonstrating that this antibody is not just an activation reporter, as it is widely used in research, but also in some manner activates LFA-1. This dual function of M24 is supported in the literature and arises from its role as an activation reporter. M24 reports activation by binding to an epitope only present in the extended open (active) state of LFA-1; following binding, M24 stabilizes this conformation resulting in a population of LFA-1 biased toward the active state ^46^.

The results from the activators, particularly the appearance of the mechano-maxima, allow us to assess how the RAD-TGT system impacts assay capability. We attributed this apparent peak to one of two sources: irreversible rupture of the TGT or the consequence of stabilizing integrins in an active state. Although possible, we do not suspect irreversible rupture because that would be accompanied by loss of cell adhesion, which was not seen. Proteins within the medium and secreted by the NK cells may also serve as compensatory adhesion points. To determine if TGTs caused the maxima, we repeated the CBR LFA-1/2 titrations on surfaces containing high-force shearing TGTs. Encouragingly, the mechano-maxima was also present, suggesting the maxima results from antibody treatment. This is further supported by reports demonstrating extended or high-concentration exposure to LFA-1 activating antibodies actually inhibits spreading, attributed to antibodies locking LFA-1 in the extended open conformation ^46^. Previously reported adhesion assays following titration of CBR LFA-1/2 see a peak of activity at the same concentration as mechano-maxima in RAD-TGT assays (**Fig S11**) ^44^.Although these adhesion assays interrogated T-cells, when paired with the results from the shearing RAD-TGTs and the paradoxical behavior of LFA-1 activators constitute a strong line of reasoning that our results were not affected by the irreversible rupture of TGTs.

To further demonstrate the ability of NATS to be used to characterize mechanotherapeutics, we sought to analyze drugs that modulate RGD-binding integrins in U-251 cells (Fig. **5C-D**). Other than providing a second therapeutic target in a markedly different system, these conditions allow us to demonstrate how the technical advances covered in this work are modular and should be used in a context-dependent manner. This is exemplified in the experimental setup for the analysis of RGD-binding integrin modulators in U-251 cells.

The first modular component is the composition of the TGTs. Since we validated no nuclease activity in U-251 cells and that they are amenable to serum-free media, we could eschew the DNA:PNA hybrid tethers and use traditional DNA duplexes. The second modular component is ligand presentation. Since we are targeting RGD-binding integrins, our previously used HUH ligand fusions are sufficient, as they already present RGD, and no crosslinking is required. Specifically, we used HUH-echistatin or HUH-fibronectin depending on the drug tested. Although both ligands are derivatives of natural RGD-presenting peptides, the fibronectin sequence used is slightly modified, containing only the N-terminal heparin-binding site fused to the 8th type III repeat with an inserted RGD sequence ^47^. Echistatin, which yields high signal, was used to identify signal loss (inhibitors), while fibronectin, with lower baseline signal, enabled detection of signal increases (activators). It should be noted that a secondary benefit of this modular design is that cost-prohibitive components, like PNA, can be avoided if conditions are amenable to cheaper alternatives.

Similar to the LFA-1 analysis, we selected three RGD-binding integrin modulating drugs: MK-0429 (an RGD-mimetic competitive inhibitor) ^48^, IDR-1002 (a synthetic innate defense response peptide reported to activate β1 integrins via PI3K-AKT signaling) ^49^, and 12G10 (a β1-specific antibody that induces integrin activation) ^50^ (Fig. **4A**). For all trials, unzipping TGTs were used. Expecting a decrease in signal, we used the high-affinity HUH-echistatin for MK-0429, and expecting the opposite, we used the lower-affinity HUH-fibronectin for IDR-1002; both ligands were used to analyze 12G10. For each condition tested (drug concentration and ligand used), an equivalent vehicle or isotype control was carried out in tandem in the same manner as the LFA-1 assay.

The RGD-binding integrin modulators generated responses similar to the LFA-1 modulators, with dose-dependent decreases of RAD-TGT signal for the inhibitor and increases for the activators. MK-0429 exhibited a robust ability to inhibit RGD-binding integrins, even when competing with the high-affinity ligand echistatin. Intriguingly, the mechano-maxima were present for the 12G10 treatment on HUH-echistatin but were absent in the IDR-1002 treatment or 12G10 on HUH-fibronectin. We initially attributed the lack of mechano-maxima in the IDR-1002 treatment to the tested concentration range, which failed to reach signal-plateauing concentrations at which the mechano-maxima would appear. However, the response of 12G10 on fibronectin TGTs challenged this interpretation. Unlike echistatin, 12G10 treatment on the fibronectin ligand appeared to behave as an inhibitor, exhibiting a dose-dependent decrease of RAD-TGT signal. This unexpected behavior is due to the integrin specificity of 12G10 combined with the composition of our HUH-fibronectin construct. The fibronectin used does not contain the 9th and 10th type III repeats; although the RGD sequence typically found in the 10th repeat is present, the PHSRN sequence found in the 9th repeat is missing. PHSRN is required for α5β1 binding ^51^, the predominant fibronectin receptor in U-251 cells ^52,53^. Thus, we reason the vexing results from fibronectin experiments stem from the specificities of 12G10 and IDR-1002 toward β1 integrins and the inability of our HUH-fibronectin construct to engage α5β1 effectively.

Intriguingly, the dose responses generated by 12G10 and IDR-1002 suggest further insights into integrin or drug function. Although relatively muted, as indicated by the lack of signal plateauing, IDR-1002 still activated integrin activity. This suggests that IDR-1002 either promotes integrin-mediated adhesion beyond β1 integrins, that conformational activation of α5β1 leads to downstream promotion of fibronectin adhesion via non-β1 integrins such as αvβ5 and αvβ6, or that IDR-1002 induces compensatory expression of the relatively low-expressed αvβ1 integrin, whose specificity toward fibronectin and epitope requirements remain ambiguous. Equally intriguing was the negative correlation between 12G10 concentration and RAD-TGT signal for probes presenting our fibronectin ligand. We postulate this response demonstrates that integrin abundance may be limited to a finite amount of activated integrins. As 12G10 concentration increases, the amount of α5β1 in the active conformation also increases, leading to sequestration of cellular resources (integrin-cytoskeleton adaptor proteins, cell surface space, etc.). This sequestration results in a reduced ability of alternative fibronectin-binding integrins to bind ligands compensatorily. Such a mechanism of integrin regulation may offer insights into how cells continually adapt to evolving surroundings. Further studies must be conducted to identify the unique behavior observed in this assay.

## Conclusion

Over the past decade, DNA-based tension sensors have emerged as a foundational tool in mechanobiology, prized for their ability to probe force transmission through specific receptors and their inherent modularity derived from DNA chemistry. In parallel with their rise, a variety of derivative tools have been developed to extend both the functional scope and technical versatility of these sensors, collectively referred to here as NATS.

Despite their broad potential, the use of NATS has remained largely confined to labs specializing in sensor development. Like many emerging technologies, widespread adoption has been limited by environmental constraints and a lack of accessible application pipelines.

In this work, we demonstrate how the integration of both established and novel strategies overcomes these barriers, expanding the environmental range and usability of NATS. More importantly, we establish NATS as a uniquely powerful platform for identifying and evaluating next generation mechanotherapeutics.

A longstanding but underreported challenge in the field has been the degradation of DNA-based sensors by nucleases. We highlight how nuclease activity can disrupt experimental workflows and show that emerging nucleic acid alternatives are compatible with high-throughput NATS . To address another major hurdle, the limited availability of ligand-DNA conjugates for researchers without synthetic chemistry resources, we developed an engineered Protein G–HUH fusion protein capable of covalently attaching Fc-containing ligands to DNA structures.

We have demonstrated that NATS enables direct, molecular-level assessment of drug-induced force transmission through therapeutic targets. Unlike traditional assays, NATS supports mechanokinetic profiling of mechanotherapeutics in primary cells, offering insight into both target engagement and force modulation. This compatibility with physiologically relevant systems paves the way for precision screening of drugs that not only bind mechanosensitive proteins, but actively modulate mechanical function, potentially on a per-patient basis.

Mechanobiology remains in its nascent stage. While mechanical forces have been implicated across nearly every aspect of cellular life, our understanding of their roles remains fragmentary. We believe the strategies outlined here, alongside the pioneering efforts of others, will help lift mechanobiology from a niche interest to an essential parameter of all life sciences.

And to that end, we offer only one warning: “Don’t underestimate the Force” ^54^.

## Methods

### Cell Culture

For non-primary cell lines (U-251 and C2C12), cells were maintained with Dulbecco’s Modified Eagle Medium (Corning, 10027CV) supplemented with 10% fetal bovine serum (FBS) (R&D Systems, S11150) and penicillin/streptomycin (Gibco, 15070063) in 10 cm tissue culture–treated dishes. Cells were regularly passaged at 80% confluency (approximately every 2–3 days) and cells were used in experiments only after passage 2 and before passage 10.

### HUH-Ligand Expression

The previously developed HUH endonuclease variants (HUH, HUH-Echi, HUH-FN) and PG-HUH were expressed and purified following previously published protocols. The PG-HUH plasmid was assembled with the PG sequence from prior works (no unnatural amino acid) appended to the N-terminus of the HUH endonuclease separated by a GGSGGS linker; a geneblock of this construct was purchased. For expression of the variants, briefly, variants were expressed in *BL21* (DE3) *Escherichia coli* (E. coli) following IPTG induction. E. coli were then lysed and protein isolated via Ni–NTA purification. The resulting protein quality was assessed via SDS–PAGE to determine if further purification via size exclusion chromatography was necessary; it was not.

### BPA Incorporation in PG-HUH

The codon for the third alanine within PG (residue #24 codon GCA) was mutated to a stop codon (TAG) via Q5 mutagenesis following the manufacturer’s (New England Biolabs) protocol and primer design tool. Following sequence confirmation of the plasmid, the plasmid was transformed into *BL21* (DE3) E. coli and plated on ampicillin agar plates. To add the BPA tRNA synthetase/tRNA pair, a single colony (from the previous sentence) was transferred to 5 mL of LB. The colony incubated at 37 °C until an OD_600_ of 0.5 was reached and was in turn cooled on ice for 15 minutes and pelleted via centrifugation at 4,000 ×*g*. The supernatant was removed, and the pellet was resuspended in cold 100 mM CaCl_2_ and incubated on ice for 30 minutes with gentle agitation every 5 minutes. Following this, centrifugation and resuspension in 200 *µ*L of 100 mM CaCl_2_ was repeated and the resuspended E. coli were transformed with the tRNA synthetase/tRNA pair plasmid, *pEVOL-pBpF* (a gift from Peter Schultz, Addgene #31190) and plated on agar containing ampicillin and chloramphenicol. The resulting colonies were grown in 2xYT media (1 L containing 16 g tryptone, 10 g yeast extract, 5 g NaCl, pH 7) containing both ampicillin and chloramphenicol. At an OD_600_ ≈ 0.3, 500 mg 4-Benzoyl-L-phenylalanine dissolved in a minimal volume of NaOH was added to the culture. At an OD_600_ ≈ 0.7, plasmid expression was induced with 0.5 mM IPTG and 0.2% (w/v) arabinose and the culture was left overnight at 18 °C. Protein was purified in the same manner as other HUH variants.

### PG Crosslinking

For all crosslinking, clPG-HUH and Fc-containing protein of interest were incubated for 20 minutes at room temperature in PBS with a 1:1.5–2 clPG-HUH Fc molar ratio. Following this, the solution was placed on ice within an AlphaThera LED Photo–Crosslinking Device (365 nm light source) for 20 minutes. For all TGT experiments, the final clPG-HUH amount was ideally 2× total DNA amount to ensure all tethers contained ligand.

### DNA Probe Preparation

For all tension gauge tethers and surface nuclease sensors, anchor and ligand strands were mixed in a 1.1:1 ratio with the fluorophore-containing strand as the lesser amount. DNA duplexes were annealed in a 1× annealing buffer (10 mM Tris pH 7.5, 50 mM NaCl, 1 mM EDTA) while PNA:DNA hybrids were annealed in water. All duplexes were heated to 98 °C for 5 min followed by cooling at room temperature for 1 h. The strand containing a quencher was used in excess to mitigate any single-stranded fluorescent strands. Duplexes to be used in TGT assays were reacted with HUH–Fusion (either no fusion, protein G, or alternative ligand) of interest in a 1:2 ratio. Reactions were performed at 37 °C for 30 min in an HUH reaction buffer (50 mM HEPES pH 8.0, 50 mM NaCl, 1 mM MnCl_2_). For the non-covalent protein–G constructs, the assembled TGT was incubated at room temperature with 2× antibody for 30 min.

### Surface Preparation

Experimental surfaces were prepared as previously described ^18^, with slight modification outlined below. Briefly, 96-well glass plates (Cellvis, P96-1.5H-N) had wells of interest washed with PBS and incubated with 80 *µ*L of 100 *µ*g/mL BSA–Biotin (Thermo Fisher, 29130) in PBS for at least 1 h at room temperature. For the SNS assay the BSA–Biotin solution also included 18.75 *µ*g/mL of fibronectin. From this point onward, 50 *µ*L excess PBS always remained in the well to prevent unwanted adsorption. Wells were rinsed 2× with cold PBS and incubated with 100 *µ*g/mL neutravidin (Thermo Fisher, 31000) in PBS for 30 min at room temperature. Wells were again rinsed 2× with PBS, and assembled probes (TGTs or SNS) were diluted to 0.1 *µ*M in PBS and 80 *µ*L was added to intended wells. Plates were sealed with adhesive aluminum foil and incubated overnight at 4 °C.

### Cell-Based Experimental Setup

For all experiments, cells were collected in 15 mL conical tubes (either following trypsinization for adherent cells or from suspension for NK cells) in appropriate culture media and washed via centrifugation at 300 ×*g* for 5 min followed by resuspension in PBS. In total, 2 washes were performed with PBS and 2 washes were performed with experimental media replacing PBS. Cell density was then assessed and adjusted to the desired cell count. For all experiments each well had 100 *µ*L of cell solution added; for RAD–TGT experiments cell density was adjusted to 150,000 cells/mL for U-251 and C2C12 cells, while NK cell density was adjusted to 500,000 cells/mL such that each well contained 15,000 or 50,000 cells, respectively. For imaging-based experiments the intended cell density was 20,000 cells/mL yielding 2,000 cells per well. Final cell amounts were determined empirically and on a cell-by-cell basis. For RAD–TGT experiments we targeted 80% confluency and for microscopy-based experiments we sought sparse cell populations with minimal cell–cell interactions.

For drug treatments, solutions containing higher concentrated drugs were prepared (either individually or via serial dilution) and sufficient volume of cell solution was added such that the final cell density and final drug concentration were as desired. For C2C12 and U-251 drug treatment, following the creation of the cell–drug solution, cells were pre-incubated at 37 °C for 30 min while NK cell treatments were pre-incubated for the same period on ice.

Either during drug incubation or prior to cell counting, plating surfaces functionalized with TGTs were rinsed 2× with PBS followed by a single rinse of appropriate experimental medium. Subsequently, 200 *µ*L experimental medium was added to acclimate cells at 37 °C in 5% CO_2_ for 1 h for all imaging and NK cell experiments and 1.5 h for RAD–TGT analysis of U-251 and C2C12 cells. Following incubation, cells were prepared for imaging (see Microscopy Methods) or for RAD–TGT assay readout (see RAD–TGT Assay).

### RAD–TGT Assay

For all RAD–TGT conditions, an equivalent negative control was prepared. For assays assessing the effects of nucleases on RAD–TGTs, every Y-27632 treatment had an equivalent volume of DMSO (vehicle control); furthermore, each treatment condition had an equivalent HUH (no ligand fusion control). For integrin modulator titrations, each drug treatment had an equivalent RAD–TGT assay set up (identical RAD–TGTs presenting identical ligands) in which cells were treated with equivalent amounts of either isotype control antibodies or DMSO (vehicle control).

To prepare for the RAD–TGT readout, cells were rinsed gently 3× with 200 *µ*L of PBS to remove culture medium and then rinsed 3× with 200 *µ*L dissociation reagent. Dissociation reagent rinses were performed quickly to mitigate any loss of cells; for weakly adhered cells only 2 rinses were performed. On the final dissociation reagent rinse, 50 *µ*L of reagent remained and an additional 25 *µ*L was added to ensure adequate volume. Cells in dissociation reagent were further incubated for 5 min or until cells started to visibly lift. Cells were resuspended in 150 *µ*L flow cytometry buffer (1 mM EDTA, 1% w/v bovine serum albumin in PBS) and gently pipetted; 200 *µ*L of the cell solution was then removed for analysis. For the first replicate of each experiment, cells were stained with propidium iodide following the manufacturer’s protocol (Cayman Chemical, 10008351) to assess viability. Viability was consistently ¿97% as the assay inherently selected for live cells. For conditions where cells struggled to adhere, trials aimed to analyze a minimum of 1,000 cell events after gating (see Flow Cytometry section).

### Peripheral Blood NK Cell Isolation and Culture

De-identified Leukocyte Reduction System (LRS) cones were obtained from the Memorial Blood Center (Saint Paul, MN). Peripheral blood mononuclear cells (PBMCs) were isolated by density gradient centrifugation using Ficoll-Paque Premium (Cytiva). NK cells were then enriched from PBMCs using the EasySep Human NK Cell Enrichment Kit (StemCell Technologies) according to the manufacturer’s protocol. NK cells were stimulated overnight in B0 media (Dulbecco’s Modified Eagle’s Medium plus Ham’s F-12 media, 2:1, supplemented with 10% heat-inactivated human AB serum, penicillin (100 U/mL), streptomycin (100 *µ*g/mL), *β*-mercaptoethanol (25 mM), ascorbic acid (20 mg/mL), and sodium selenite (5 ng/mL)) with IL-15 (10 ng/mL; National Cancer Institute) at a density of 3 × 10^6^ cells/mL. Following overnight stimulation, NK cells were counted, harvested, and washed with B0 to readjust the cell concentration to 500,000 cells/mL and the cytokine concentration to 1 ng/mL IL-15 for subsequent assays.

### Flow Cytometry

For all flow cytometry experiments, cells were resuspended in flow buffer (1 mM EDTA, 1% w/v bovine serum albumin in PBS) and stored on ice from sample collection to data analysis. For non-RAD–TGT experiments, cell viability was assessed with propidium iodide following the manufacturer’s protocol (Cayman Chemical, 10008351). To create a positive viability control, unstained samples were incubated at 65 °C to generate a verified dead population and stained with propidium iodide.

For all flow cytometry experiments (barring RAD–TGTs), a minimum of 10,000 cells per final gated population were collected. Gating was performed in the following sequence (see Supplementary Figure X): cells were first gated by size via FSC-A vs. SSC-A dot plots of the sample input. Single cells were gated using FSC-A vs. FSC-H plots. Live cells were gated by propidium iodide signal vs. FSC-A. For some channels (e.g. CY5 intensity), a population of zero fluorescent intensity (below negative control) cells was observed, attributed to a technical artifact of the BD Accuri C6 Plus. In these cases, a vertical gate was set at a low fluorescence intensity of 10 units (below negative control) and cells exceeding the gate were selected.

Following collection, all samples were re-gated using FlowJo version 10.10. Compensation controls were performed with single-stained controls for all fluorophores used. Processed data were exported for further analysis.

### Microscopy

For all microscopy-based experiments, cells were first fixed with 4% paraformaldehyde for 10 min and washed gently 3–5× with cold PBS. If intracellular staining was needed, cells were permeabilized via 10 min incubation in permeabilization buffer (3% BSA, 0.25% Triton X-100 in PBS) and washed at least 3× with PBS. Cells were stained with phalloidin and/or DAPI following manufacturers’ recommendations. Cells were imaged with an Olympus IX83 motorized inverted fluorescence microscope. For qualitative image creation, ImageJ was used with the Dotted Line plug-in. For cell outlines, phalloidin-stained images were outlined and the outline was superimposed via the Dotted Line plug-in onto images.

### Image Quantification

For quantitative analysis of microscopy images, CellProfiler was used. Briefly, for cell spreading area images, the phalloidin channel (actin cytoskeleton) and DAPI channel (nuclei) were used. Phalloidin images were thresholded to identify cell outlines. Outlines containing more than one nucleus or touching the edge of the image were excluded. The resulting isolated single cells then had their total area measured. For nuclease activity, images contained three channels: phalloidin, DAPI, and SNS (nuclease activity). Cell outlines were identified by thresholding the phalloidin channel; SNS activity extending beyond the cell outline was captured by thresholding the SNS channel. Regions of thresholded SNS activity overlapping cell outlines were demarcated as “under cell” and all other areas as background. The median intensity of each under-cell object and background were recorded. The average of all under-cell median intensities for serum-free U-251 cells was used to normalize all data measurements within each replicate. All under-cell medians and background medians were divided by this average to normalize data.

### RAD–TGT Quantification

Following data collection and processing (see Flow Cytometry), relative RAD–TGT signal was quantified via a percent-past-control gate or fold-change metric (as noted in figure legends). Main text figures used percent-past-control gate; supplement figures used fold change. Each experimental condition had a respective negative control. For drug titrations, every concentration had an equivalent negative control well with identical RAD–TGTs but cells treated with vehicle or isotype controls. For single-drug effects (Y-27632), negative controls were wells with RAD–TGTs reacted with HUH (no ligand fusion) instead of HUH-Ligand.

To quantify percent-past-control, a vertical gate at the 99th percentile of each negative control histogram was applied to each experimental population and the percentage exceeding that gate was recorded. For fold change, the median intensity of the negative control was recorded and each experimental median was divided by that value.

For drug titration values, data were normalized to a 0–100 scale. For inhibitors, the no-treatment concentration was set as 100 and the concentration generating the lowest signal as 0. For activators, the no-treatment concentration was set as 0 and the highest-signal concentration as 100. Normalization was performed per replicate prior to comparisons. Curves were fitted using GraphPad Prism (v10.3.0) built-in [Inhibitor] vs. response (three parameter) and [Agonist] vs. response (three parameter) functions.

### Graphing and Statistics

All statistical analyses and graphs were generated with GraphPad Prism (v10.3.0) as noted in figure legends. For superplot ^55^, only replicate means (large opaque symbols) were used for statistical analysis. All figures were finalized in Inkscape (v1.4).

## Notes

### Competing Interest Statement

The authors have declared no competing interest.

