## Supplementary information for "A Generalizable Tension Sensor Platform for Mechanotherapeutic Discovery"

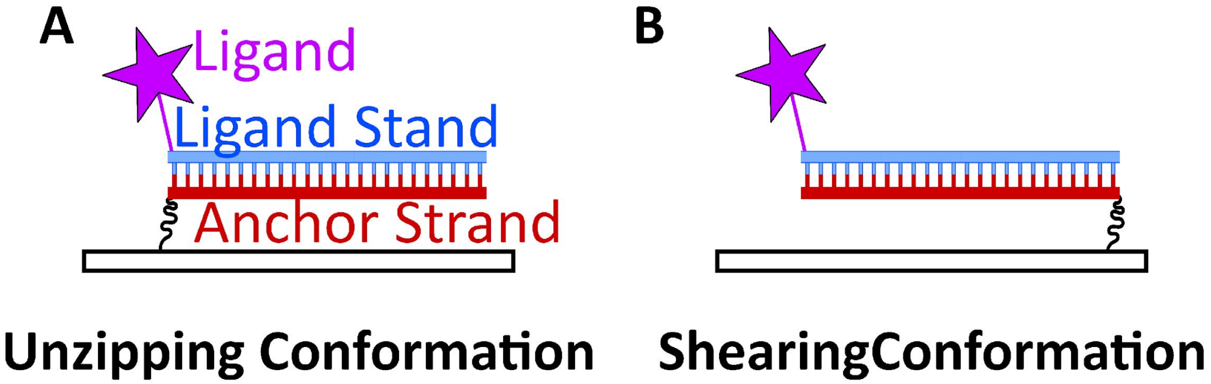


Figure S1: Tension gauge tether (TGT) are DNA duplexes in which one strand possess a ligand for a receptor of interest (the ligand strand) and one strand possesses a fixation moiety (anchor strand). A) When the ligand and fixation moieties are on proximal termini the TGT is in unzipping conformation. The unzipping conformation requires the least force to rupture. B) If the ligand and fixation moieties are on distal termini the duplex is in the shearing conformation requiring greater force to rupture. Intermediate forces are achieved by altering relative position of the fixation moiety to the ligand.

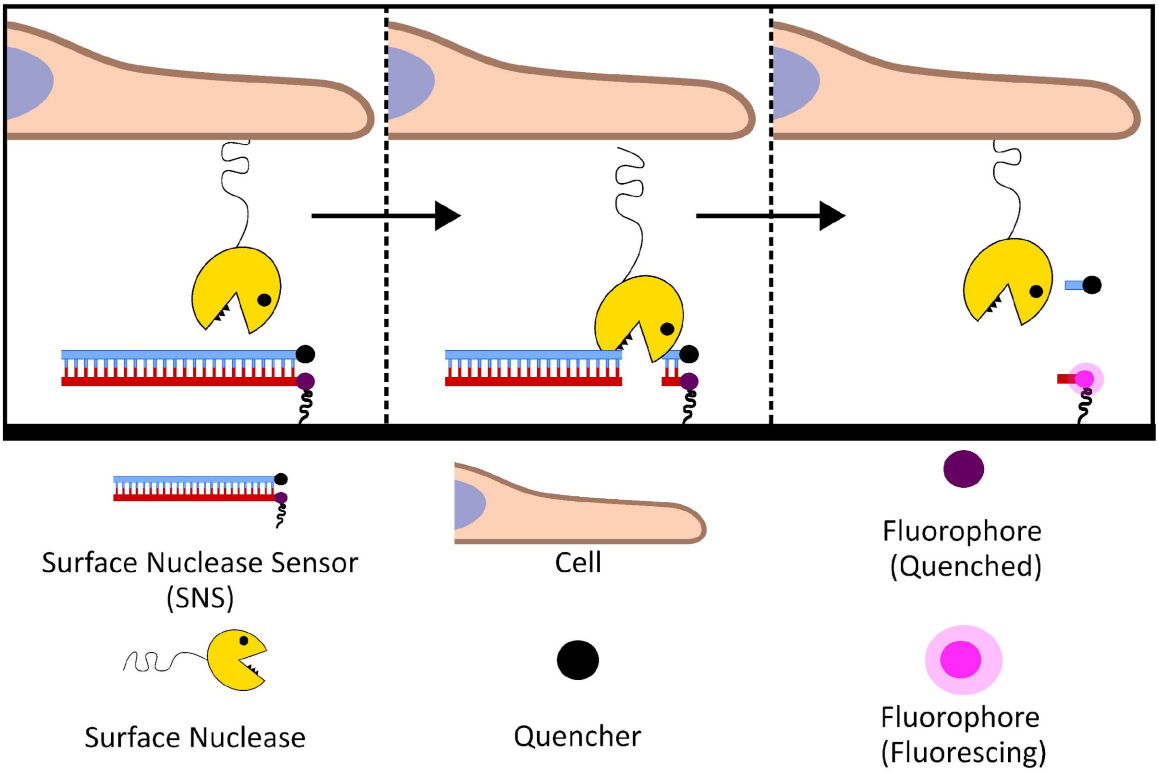


Figure S2: Cartoon of principle behind surface nuclease sensors (SNS). SNS are similar to TGTs but lack the ligand. A fluorophore is near the fixation moiety while a quencher is on the opposing strand. If nucleases cleave the duplex the duplex dissociates resulting in the fluorophore quencher pair separating and a fluorescent signal on the surface where the nuclease cleaved.

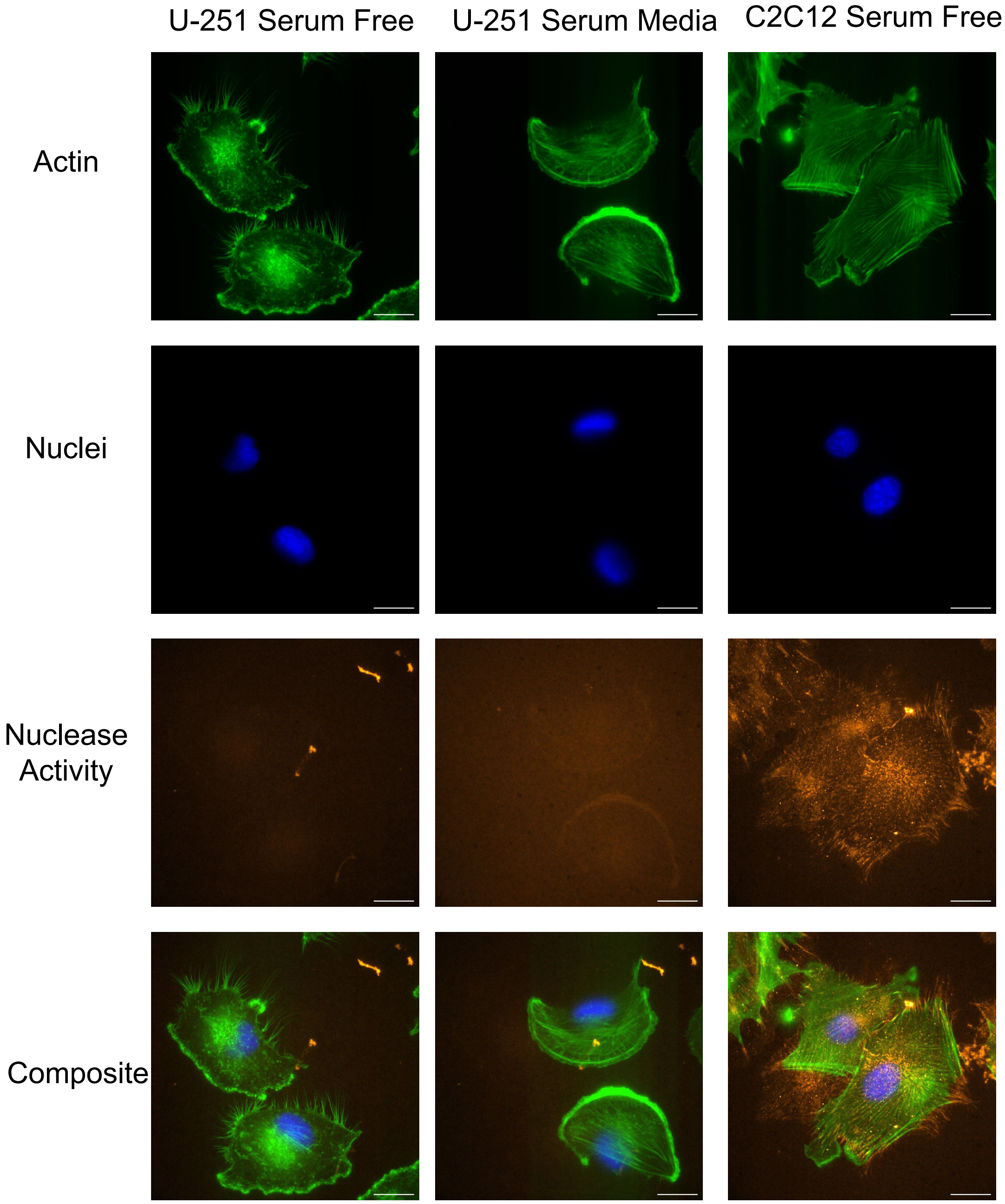


Figure S3: Representative images of cells assessed for nuclease activity. All three conditions from figure 1 with all channels imaged to assess nuclease activity: phalloidin for actin cytoskeleton, DAPI to image nuclei, SNS to measure nuclease activity, and a composite of all channels overlaid. Scale = 10 μM.

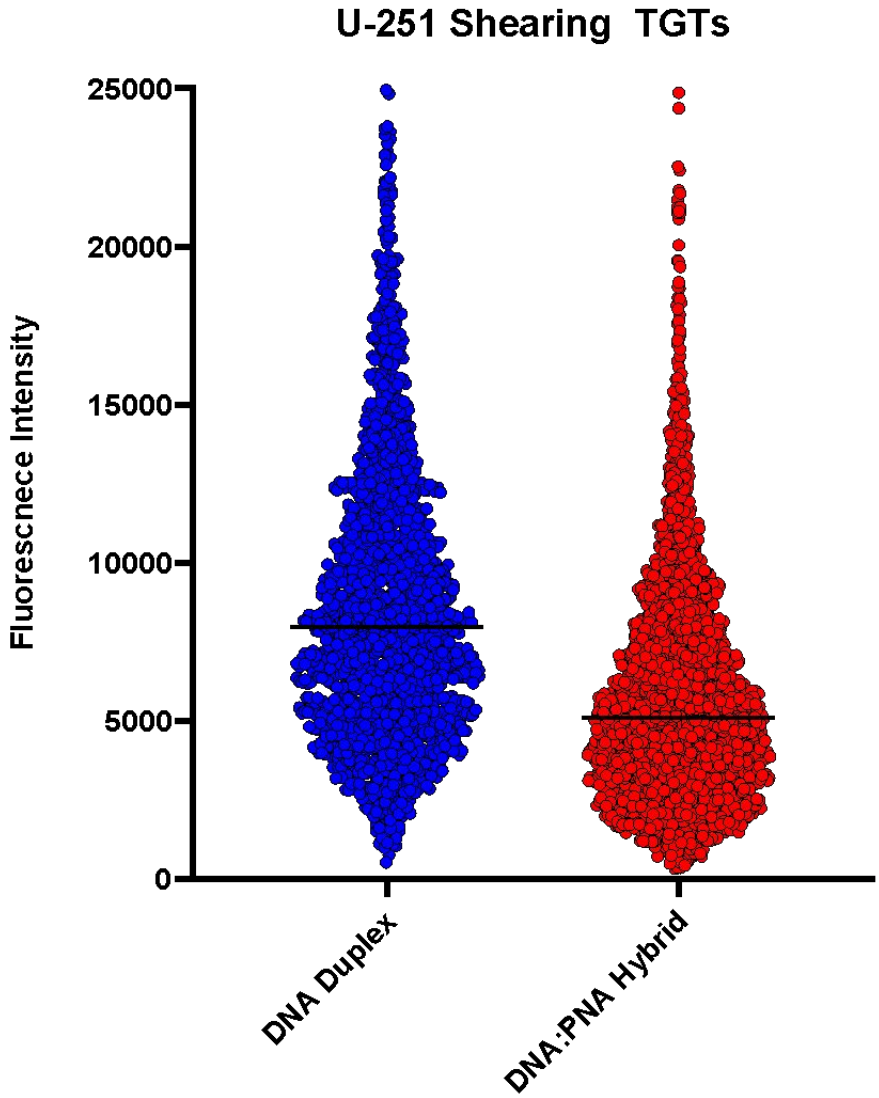


Figure S4: Representative graph of fluorescent intensity generated by U-251 cells on shearing RAD-TGTs with echistatin as ligand on either tethers composed of DNA (blue) or DNA:PNA hybrid (red). Horizontal black line is median intensity.

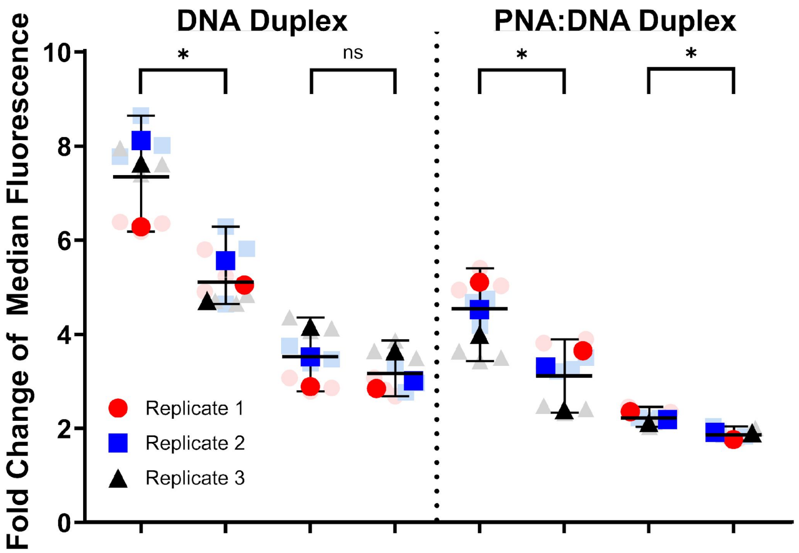


| Cell Line | U | U | C | C | U | U | C | C |
|---|---|---|---|---|---|---|---|---|
| Y-27632 | - | + | - | + | - | + | - | + |

Figure S5: Quantification of Y-27632 effect on low nuclease cells (U-251) or high nuclease cells (C2C12) on TGTs composed of DNA or DNA:PNA hybrids using fold change of median intensity. Identical data set and notation to figure 2C, quantification differs from 2C utilizing fold change of experimental median relative to the median of each respective negative control. Ns, not significantly different. *$p<0.05$, unpaired students t-test.

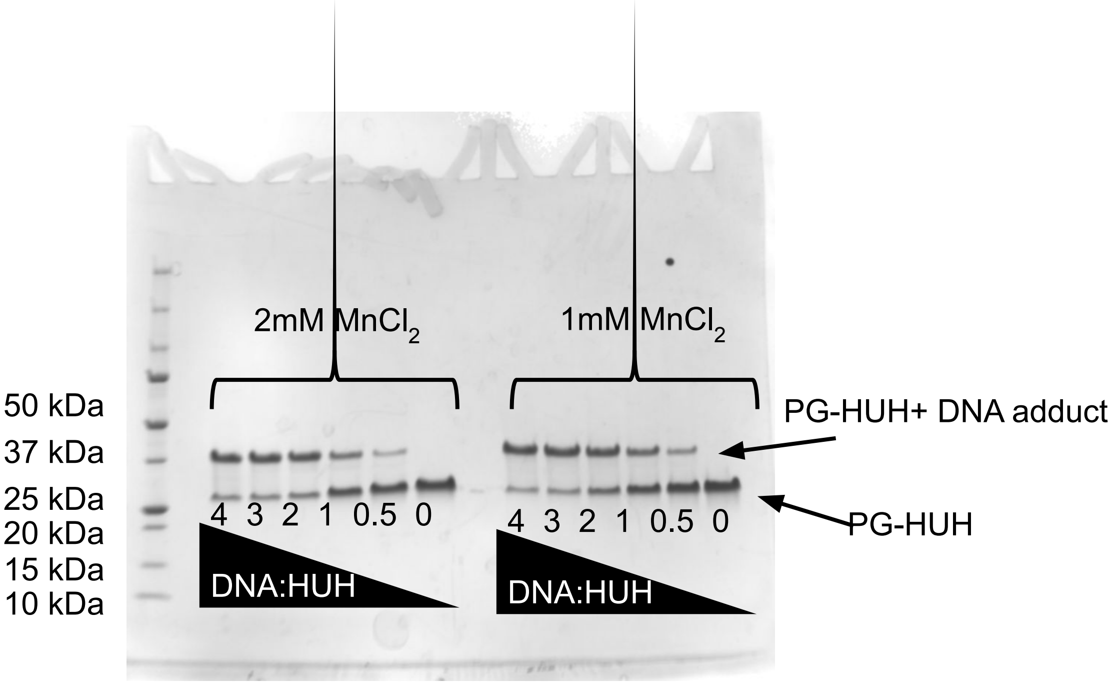


Figure S6: Assessing ability of PG-HUH to form covalent adduct with DNA. HUH Oligo reactions were prepared in either 4,3,2,1,0.5,or 0:1 molar ratios of DNA to PG-HUH respectively in the presence of either 2mM $MnCl_2$ or 1 mM $MnCl_2$. Construct retained HUH activity.

A

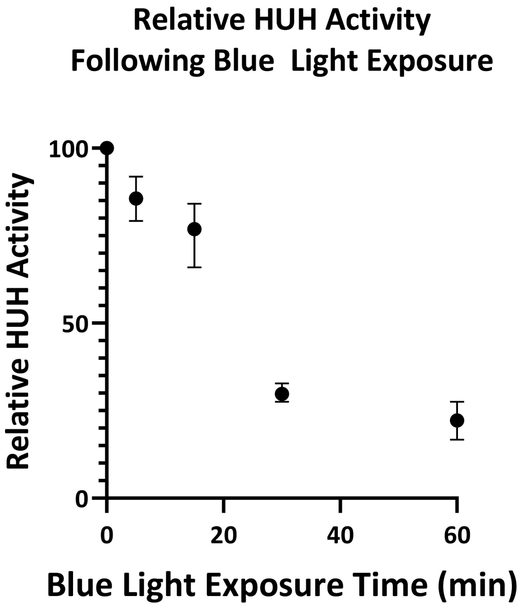


B

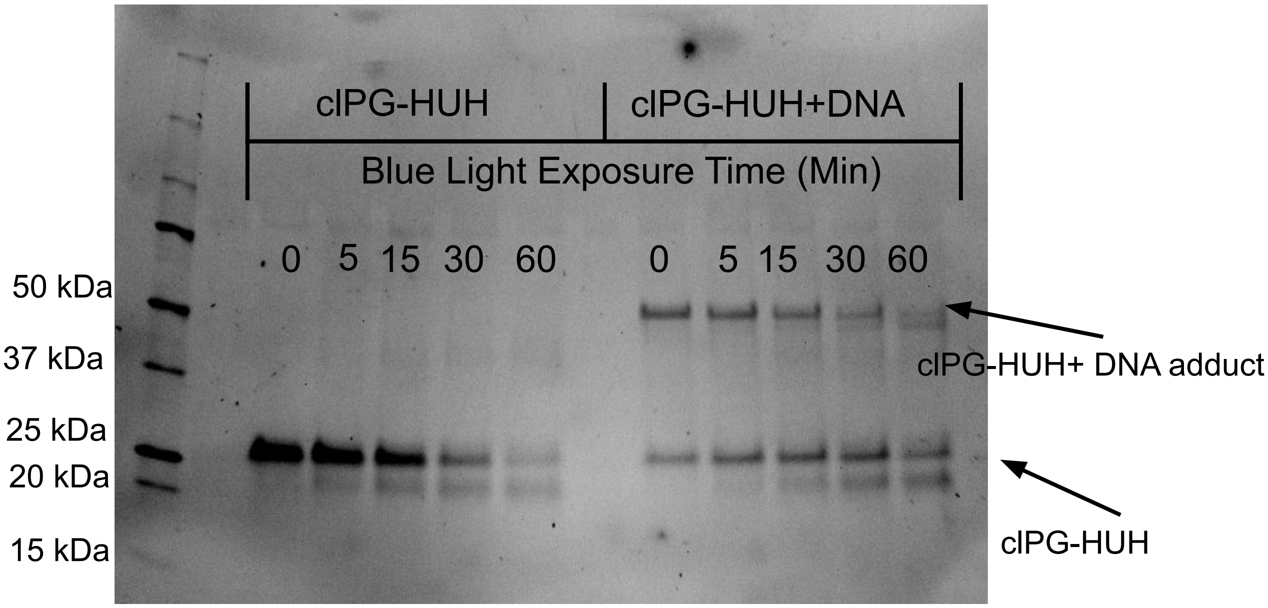


Figure S7: DNA Binding Activity of clPG-HUH is dependent on length of blue light exposure. A) clPG-HUH was exposed to blue light for either 0,5,15,30 or 60 minutes. Following exposure time, samples were mixed with DNA in a 1.5:1 ratio (HUH to DNA) and an HUH reaction was carried out, samples with or without DNA were analyzed via SDS-PAGE. Activity was normalized relative to 0-minute exposure time, errors bars represent normalized range of 3 replicates 3. B) Representative image of SDS-PAGE results

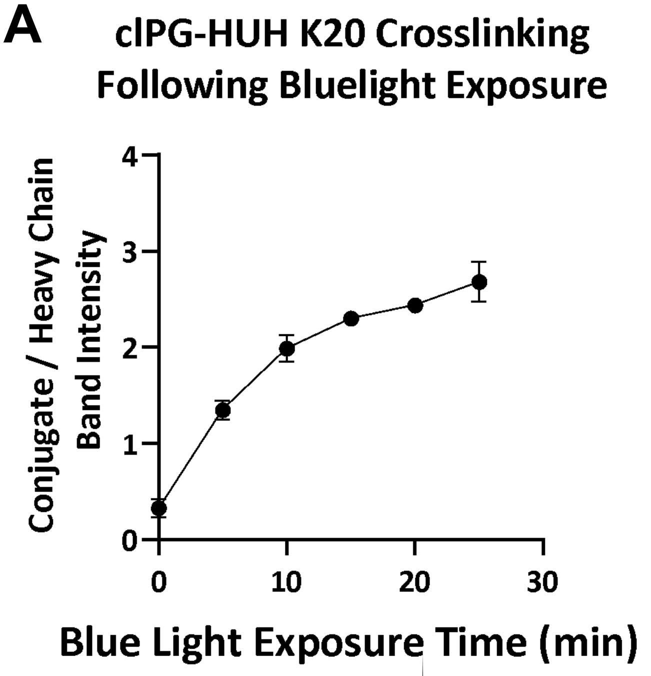


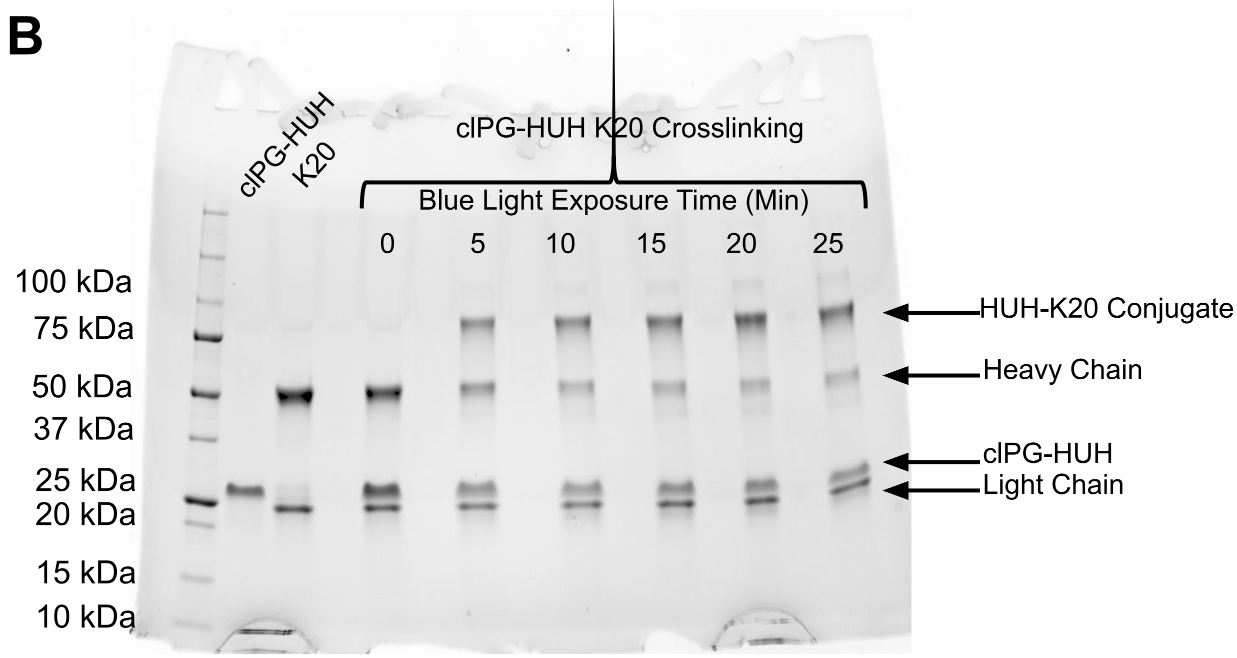


Figure S8: cIPG-HUH antibody conjugation is exposure time dependent. A) cIPG-HUH antibody mixtures were exposed to blue light for either 0,5,10,15,20, or 30 minutes. Samples contained a molar ratio of 2:1 cIPG-HUH to antibody (K20). Band intensity for HUH-K10 conjugate and heavy chain were measured, at 0 min location of theoretical conjugate location was measured. Errors bars represent normalized range of 2 replicates . B) Representative image of SDS-PAGE results

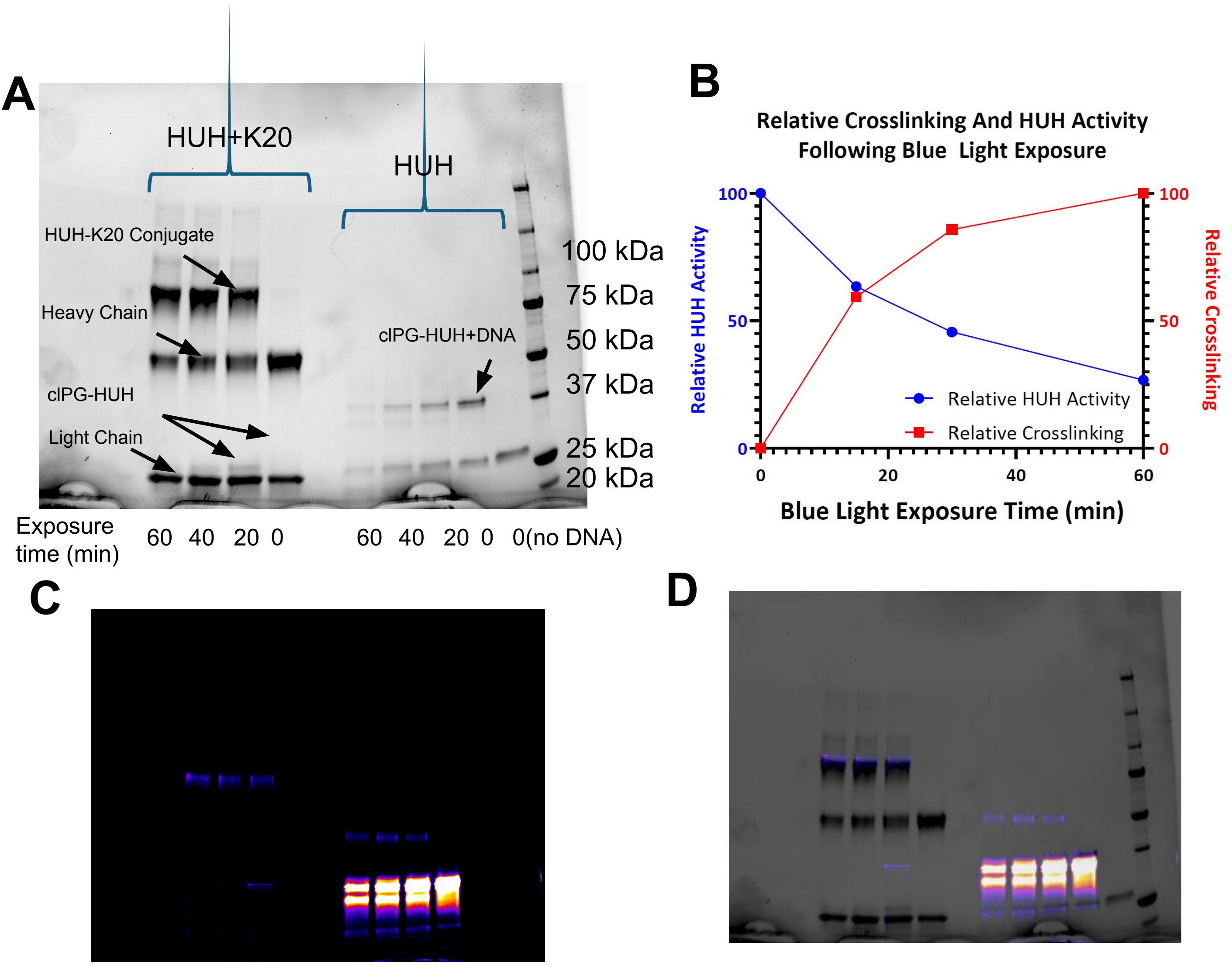


Figure S9: At either a 1:0 or 1:2 molar ratio of clPG-HUH and antibody(K20) were reacted and exposed to blue light for either 0,20,40, or 60 minutes. Following this the resulting products were reacted with 2x excess fluorescently labelled oligo. A) Resulting samples were analyzed via SDS-PAGE. B) Relative HUH activity and crosslinking were determined by quantifying band intensity. For HUH activity clPG-HUH + DNA conjugate band intensity was divided by intensity of clPG-HUH band intensity, values were normalized by setting the highest activity lane to 100, and the no oligo control to 0 (intensity of clPG-HUH+DNA conjugate for the no oligo control was region intensity of theoretical band location). For relative crosslinking HUH-K20 conjugate band intensity was divided by intensity of heavy chain band intensity, values were normalized by setting the highest activity lane to 100, and the 0 min control to 0 (intensity of HUH-K20 conjugate for the 0 min control was region intensity of theoretical band location.) C) Identical gel to A but imaged via fluorescent imager to detect oligo. D) Overlay of figure A and C to demonstrate HUH activity across conditions. N = 1

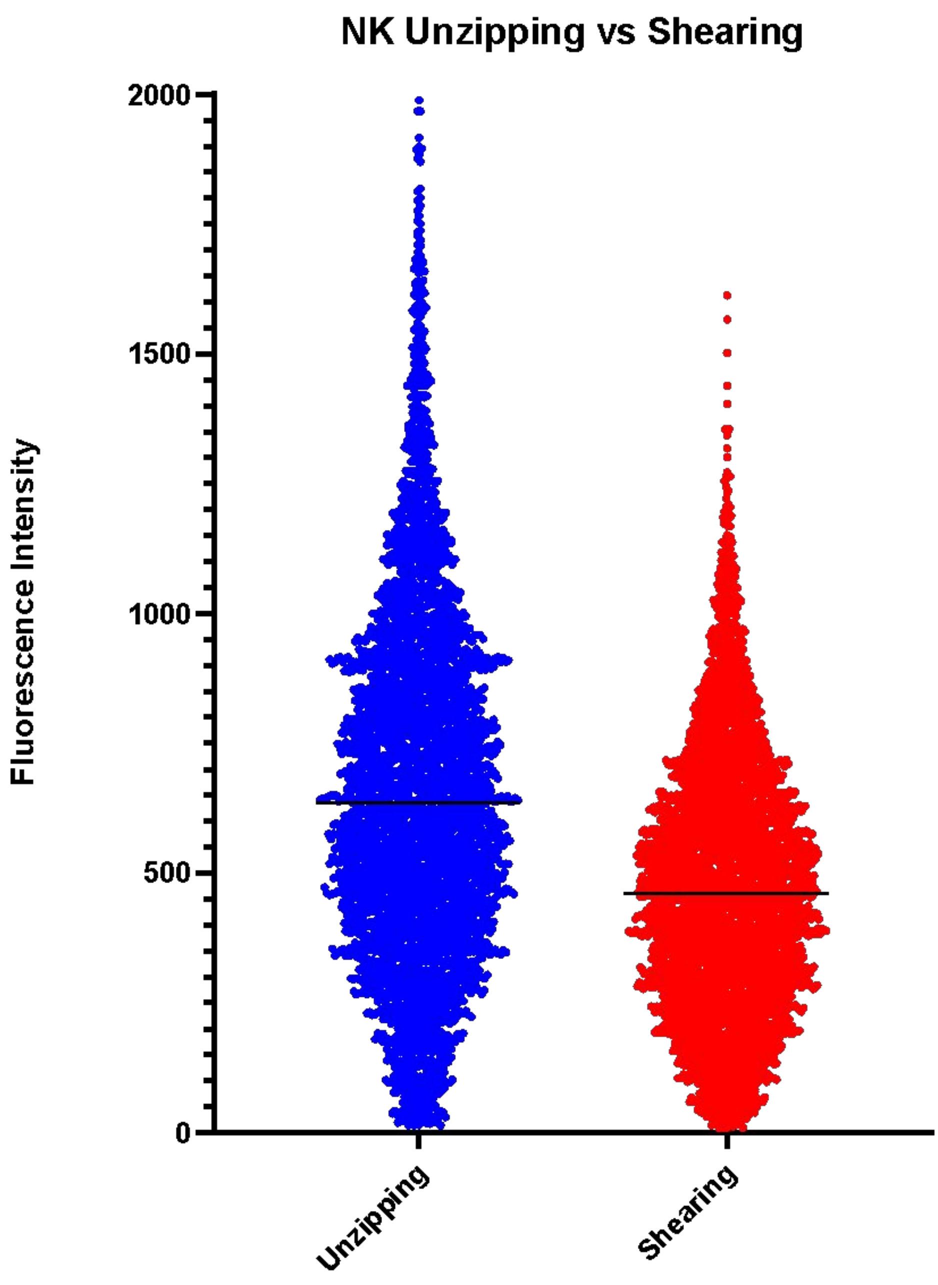


Figure S10: Representative graph of fluorescent intensity generated by NK cells on unzipping (blue) or shearing (red) RAD-TGTs with ICAM-1. Horizontal black line is median intensity.

A

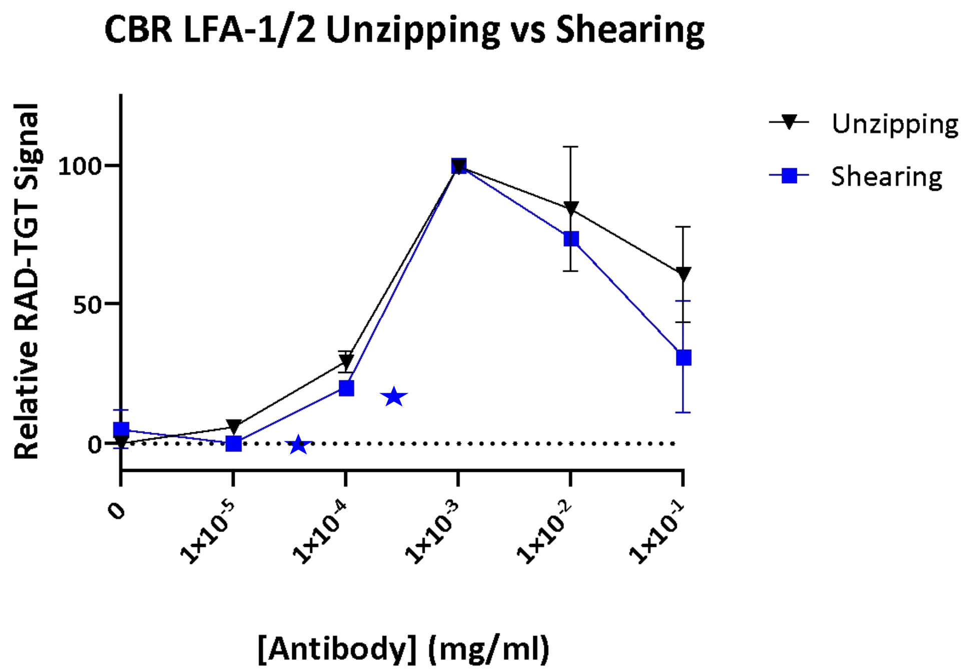


B

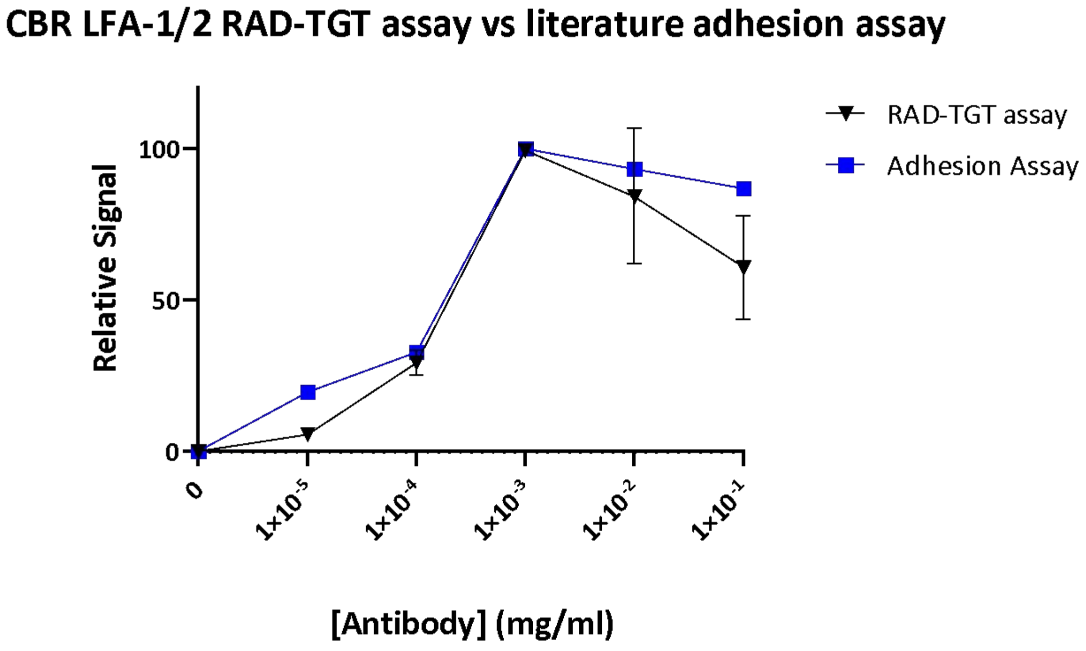


Figure S11: Data from figure 4A from NK cells treated with CBR LFA-1/2 on unzipping RAD-TGTs overlayed with alternative readouts. A) 4A data (black) overlayed with experimental data from identical experiment utilizing shearing (high requisite force) TGTs (blue) as opposed to unzipping. Shearing data is the mean of 2 independent experiments scaled from 0 to 100, except for points marked with a blue star in which only 1 data point is captured. At least 2000 cells analyzed for each replicate. Error bars represent standard deviation of replicates. B) 4A data (black) overlayed with adhesion assay data of lymphocytes treated with a CBR LFA-1/2 titration from figure 10B from Petruzzelli *et al.* 1995 (blue).
